# MEMO: Mass Spectrometry-based Sample Vectorization to Explore Chemodiverse Datasets

**DOI:** 10.1101/2021.12.24.474089

**Authors:** Arnaud Gaudry, Florian Huber, Louis-Félix Nothias, Sylvian Cretton, Marcel Kaiser, Jean-Luc Wolfender, Pierre-Marie Allard

## Abstract

In natural products research, chemodiverse extracts coming from multiple organisms are explored for novel bioactive molecules, sometimes over extended periods. Samples are usually analyzed by liquid chromatography coupled with fragmentation mass spectrometry to acquire informative mass spectral ensembles. Such data is then exploited to establish relationships among analytes or samples (e.g. via molecular networking) and annotate metabolites. However, the comparison of samples profiled in different batches is challenging with current metabolomics methods. Indeed, the experimental variation - changes in chromatographical or mass spectrometric conditions - often hinders the direct comparison of the profiled samples. Here we introduce MEMO - **M**S2 Bas**E**d Sa**M**ple Vect**O**rization - a method allowing to cluster large amounts of chemodiverse samples based on their LC-MS/MS profiles in a retention time agnostic manner. This method is particularly suited for heterogeneous and chemodiverse sample sets. MEMO demonstrated similar clustering performance as state-of-the-art metrics taking into account fragmentation spectra. More importantly, such performance was achieved without the requirement of a prior feature alignment step and in a significantly shorter computational time. MEMO thus allows the comparison of vast ensembles of samples, even when analyzed over long periods of time, and on different chromatographic or mass spectrometry platforms. This new addition to the computational metabolomics toolbox should drastically expand the scope of large-scale comparative analysis.

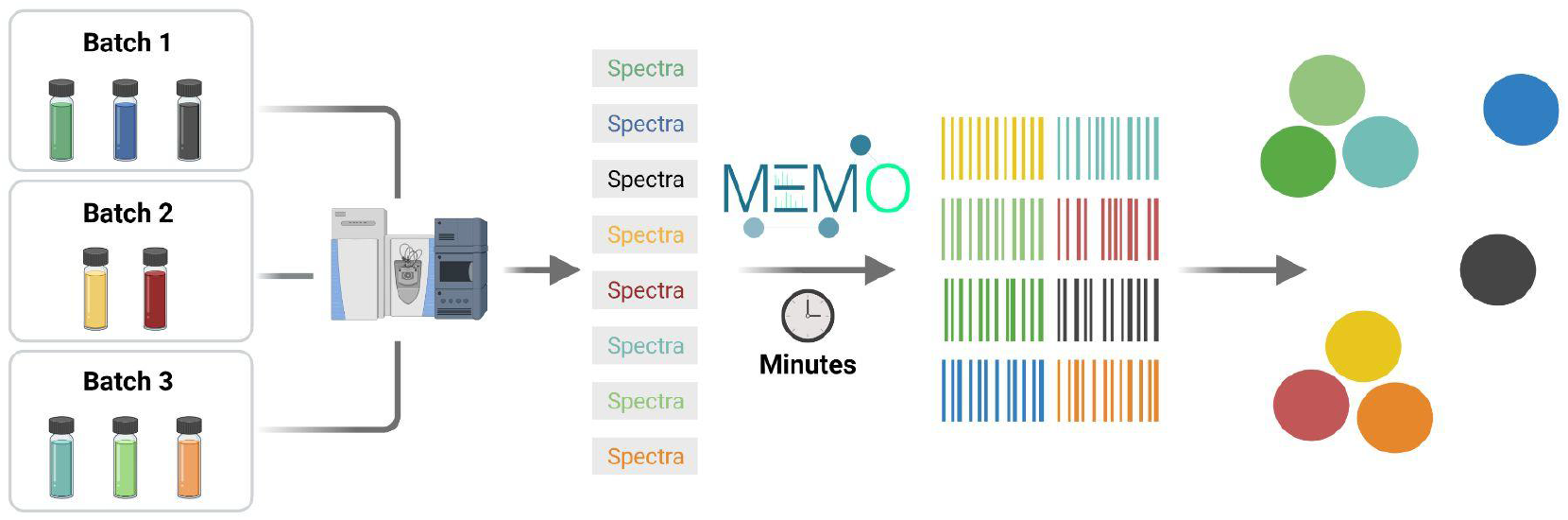

## Introduction

Mass spectrometry (MS) has been and remains the workhorse of the natural products (NP) community. It is widely used to profile complex extracts, especially using untargeted tandem mass spectrometry experiments hyphenated with liquid chromatography (LC-MS/MS) (Wolfender et al., 2019). Following untargeted analysis in data dependent acquisition (DDA), the samples data are usually processed using different tools such as MZmine, MS-Dial, or XCMS to detect features (i.e., *m/z* values and their respective retention time (RT)) and link those to their associated MS/MS spectrum (Smith et al., 2006; Pluskal et al., 2010; Tsugawa et al., 2015). The resulting feature lists and fragmentation spectra are then used to perform multivariate analysis for comparing different samples and/or Molecular Networking (MN) along with spectral annotation (Allard et al., 2016; Wang et al., 2016; Rutz et al., 2019). Despite recent progress, current sample clustering approaches face severe limitations, in particular when comparing chemically diverse samples in large datasets, as is often the case in NP research.

Typically, metabolomics comparison is carried out on tables constituted by features intensities across samples. An issue frequently observed when comparing such large datasets - often acquired over days, months or even years - is the lack of reproducibility both on the LC side with retention time shifts and on the MS detection side with variation in sensitivity and accuracy of the mass spectrometer (Dunn et al., 2012; Arens et al., 2015). Strategies have been devised to reduce and balance these so-called “batch-effects”, but are often not adapted to heterogeneous NP extracts datasets where the feature overlap is weak due to the large chemodiversity of the profiled matrices (Wehrens et al., 2016). Classical multivariate analyses such as principal component analysis (PCA) or principal coordinates analysis (PCoA), but also other dimensionality reduction techniques such as uniform manifold approximation and projection (UMAP) or tree map (TMAP), are sensitive to these effects, complicating the clustering of samples (Leek et al., 2010; McInnes et al., 2018; Probst and Reymond, 2020). Moreover, while numerous tools are designed to treat data originating from a single experiment, few techniques allow to efficiently compare samples analyzed in multiple experiments (i.e. different instrument, method and/or analysis batches). As mentioned by Jarmusch and co-authors, such data-mining techniques at the repository scale level are needed to compare data originating from different experiments, even in a single laboratory, and allow optimal data reuse (Jarmusch et al., 2021).

To reflect the chemical diversity of heterogeneous sample collections, while tackling batch-effects, several methods have been introduced that take into account the relationships among analytes (notably via their MS/MS spectrum or chemical structure). This assumption of relatedness among metabolites stands in opposition with the classical multivariate analyses which consider each feature as an independent variable. The first of these methods considering relationships among variables was developed in 2017 by Sedio *et al*, and called the chemical structural and compositional similarity (CSCS). The CSCS exploits the cosine similarity between MS/MS spectra in combination with their relative abundance in the samples to compute a distance matrix (Sedio et al., 2017). The recently developed Qemistree approach uses predicted molecular fingerprints from features’ MS/MS spectra thanks to SIRIUS and CSI:FingerID (Duhrkop et al., 2015, 2019), to generate a “tree” of metabolite relationships (Tripathi et al., 2021) that is further used to compute a UniFrac distance among samples (Lozupone and Knight, 2005). UniFrac was initially developed to compare microbial communities and was adapted to metabolomics data by Junker in 2018 with, in this case, the enzymatic proximity of the metabolites being used for the tree generation (Junker, 2018). While improving classical sample classification approaches, these methods are computationally expensive, particularly when applied on large datasets, as pairwise comparison of all spectra or fingerprints is necessary. Moreover, an alignment step yielding a feature table (giving the occurrence and/or intensity of each feature in each sample) is typically required to compare samples using these methods. Such alignments are usually based on retention time (RT) comparison and thus imply the use of comparable chromatographic methods to profile the studied samples in order to obtain a set of unique features aligned across the studied samples.

Here we introduce MEMO - **M**S2 Bas**E**d Sa**M**ple Vect**O**rization - a method allowing the efficient comparison of large and heterogeneous sample ensembles based on their LC-MS/MS profiles. The first step of the MEMO approach consists in extracting the fragment ions and neutral losses from each binned MS/MS spectra of the detected features (m/z @ RT) into so-called *documents*. Then, for a given sample, all created documents are aggregated based on word occurrences to constitute a fingerprint, hereafter defined as a *MEMO-vector*. Since words are purely based on mass spectrometry information, the MEMO vectors resume the full spectral diversity of a sample in a RT-agnostic fashion. These MEMO vectors can then be aligned and used for classical multivariate analysis (PCA, PCoA) and/or other dimensionality reduction approaches such as UMAP or TMAP (Probst and Reymond 2020; McInnes et al. 2018), allowing the efficient calculation and visualization of spectral relationships within large cohorts of samples. The MEMO strategy exploits the advantages of LC, namely its separative power (thus simplifying the chemical complexity of the analyzed sample and allowing isomerism resolving) while avoiding the caveats of a RT based alignment since the MEMO vectors only contain mass spectrometric information. Hereafter, we first detail the MEMO process and its implementation. We then present the comparison of the MEMO approach to currently existing state-of-the-art MS fragmentation-based sample organization approaches, namely CSCS and Qemistree using a previously published evaluation dataset (Tripathi et al., 2021). Finally, to investigate and showcase the scalability of the MEMO approach, we present the results of its application on a large and heterogeneous plant extracts dataset (1,600 extracts) which is currently exploited in our laboratory to search for novel bioactive molecules.

## Material and Methods

### Evaluation dataset

#### Description of the evaluation dataset

For the evaluation of the MEMO method, we used a dataset from the Qemistree method that consists of four « parent » chemo-diverse samples (one plasma, one tomato and two feces samples), binary mixtures (in different proportions) and quaternary mixtures of these four « parent samples ». This resulted in 27 samples that were acquired in triplicates using two different RP-UHPLC methods simulating a Retention Time (RT) shift and thus generating a strong artificial batch effect (named C18 and C18 RT-shift) and two different Mass Spectrometers (Q-Exactive Orbitrap (QE) and Maxis II Q-TOF (Q-ToF)). For more details on this dataset, see (Tripathi et al., 2021).

#### LC-MS/MS analysis and data-processing

For details on LC-MS/MS analysis and data processing, see Tripathi *et al* 2021 (Tripathi et al., 2021). Data and methods have been deposited on the GNPS/MASSive repository under accession number [MSV000083306](https://massive.ucsd.edu/ProteoSAFe/dataset.jsp?task=c5312f8ecb7d45feb35a723ba68874d2). The GNPS Feature-Based Molecular Networking (FBMN) job is available [here](https://gnps.ucsd.edu/ProteoSAFe/status.jsp?task=044e981ff0d84246ae5c91ef3db643a8) and the GNPS Qemistree job is available [here](https://gnps.ucsd.edu/ProteoSAFe/status.jsp?task=8ca56d6e33bc4106b46ba5e3510c91cb). The spectra file and the feature table used for classical PCoA, MEMO and CSCS were the one used for the FBMN job.

#### MEMO analysis

Spectra were processed using memo_ms v0.1.3 ([available on pypi](https://pypi.org/project/memo-ms/0.1.3/)) and the following parameters for spectra import were used: min_relative_intensity: 0.01, max_relative_intensity: 1, min_peaks_required: 10, losses_from: 10, losses_to: 200, n_decimals: 2. No filtering was applied on the resulting MEMO matrix. For the comparison between QE and Q-ToF data, matrices were normalized sample-wise. The distance matrices were computed using pdist function (‘braycurtis’ metric) from scipy package (1.7.1) and scikit-bio (0.5.6) was used for PCoA computation on the resulting distance matrix (Virtanen et al., 2020).

#### CSCS analysis

The CSCS analysis was run using the Qiime 2 (https://github.com/madeleineernst/q2-cscs) implementation of the chemical structural and compositional similarity metric initially developed by (Sedio et al., 2017). CSCS was calculated using the Qiime2 (Caporaso et al., 2010) CSCS implementation (https://github.com/madeleineernst/q2-cscs). Qiime2 release version 2021.2 was employed. The Qemistree set feature table was appropriately formatted as Qiime FeatureTable[Frequency] artifact using the following (https://github.com/mandelbrot-project/memo_publication_examples/blob/main/02_qemistree/qemistree_formatter_for_cscs.py). The parameters to launch the CSCS were as follows: minimal cosine score between two features to be included (--p-cosine-threshold): **0.7**; perform Total Ion Current Normalization (TIC) on the feature table (--p-normalization): **yes**; parallel processes to run (--p-cpus): **40**; weight CSCS by feature intensity (--p-weighted / --p-no-weighted): **yes** and **no**, respectively, for the weighted and unweighted sets. To run CSCS with the unweighted option, the following minimally modified script was used (https://github.com/madeleineernst/q2-cscs/pull/3).

#### Qemistree analysis

To generate the Qemistree distance matrix from the GNPS Qemistree task [output](https://gnps.ucsd.edu/ProteoSAFe/status.jsp?task=8ca56d6e33bc4106b46ba5e3510c91cb), the code from the [Evaluation Dataset Notebook](https://github.com/knightlab-analyses/qemistree-analyses/blob/master/Evaluation-Dataset-Analyses.ipynb) of Tripathi et. al, available on Github, was used with slight adaptations to allow export of the distance matrix (the modified script is available https://github.com/mandelbrot-project/memo_publication_examples/blob/main/02_qemistree/qemistree_dm_generation.py). It is to be noticed that for the Qemistree workflow, only features with an m/z < 600 and a Zodiac score > 0.98 were used for fingerprinting and downstream analysis, resulting in 3776 of 7032 features being considered (Ludwig et al., 2020; Tripathi et al., 2021).

### Plant extract dataset & associated bioactivities

#### Description of the plant extract dataset

A plant extract library under investigation for drug discovery purposes in our lab was profiled by untargeted LC-MS. This dataset is in fact a random and heterogeneous subset of 1,600 extracts of the Pierre Fabre Research Institute Library (European Commission, 2020). This collection is constituted by EtOAc extracts, thereby yielding mostly specialized metabolites of intermediate polarity. Raw and processed data have been deposited on the GNPS/MASSive repository accession number MSV000087728.

#### LC-MS/MS analysis

Chromatographic separation was performed on a Waters Acquity UPLC system interfaced to a Q-Exactive Focus mass spectrometer (Thermo Scientific, Bremen, Germany), using a heated electrospray ionization (HESI-II) source. Thermo Scientific Xcalibur 3.1 software was used for instrument control. The LC conditions were as follows: column, Waters BEH C18 50 × 2.1 mm, 1.7 μm; mobile phase, (A) water with 0.1% formic acid; (B) acetonitrile with 0.1% formic acid; flow rate, 600 μl·min-1; injection volume, 6 μl; gradient, linear gradient of 5-100% B over 7 min and isocratic at 100% B for 1 min. The optimized HESI-II parameters were as follows: source voltage, 3.5 kV (pos); sheath gas flow rate (N2), 55 units; auxiliary gas flow rate, 15 units; spare gas flow rate, 3.0; capillary temperature, 350.00°C, S-Lens RF Level, 45. The mass analyzer was calibrated using a mixture of caffeine, methionine–arginine–phenylalanine–alanine–acetate (MRFA), sodium dodecyl sulfate, sodium taurocholate, and Ultramark 1621 in an acetonitrile/methanol/water solution containing 1% formic acid by direct injection. The data-dependent MS/MS events were performed on the three most intense ions detected in full scan MS (Top3 experiment). The MS/MS isolation window width was 1 Da, and the stepped normalized collision energy (NCE) was set to 15, 30 and 45 units. In data-dependent MS/MS experiments, full scans were acquired at a resolution of 35,000 FWHM (at m/z 200) and MS/MS scans at 17,500 FWHM both with an automatically determined maximum injection time. After being acquired in a MS/MS scan, precursor ions were placed in a dynamic exclusion list for 2.0 s.

#### Data-processing

The MS data were converted from .RAW (Thermo) standard data format to .mzXML format using the MSConvert software, part of the ProteoWizard package (Chambers et al., 2012). The converted files were treated using the MZmine software suite v. 2.53 (Pluskal et al., 2010). The parameters were adjusted as follows: the centroid mass detector was used for mass detection with the noise level set to 1.0E4 for MS level set to 1, and to 0 for MS level set to 2. The ADAP chromatogram builder was used and set to a minimum group size of scans of 5, minimum group intensity threshold of 1.0E4, minimum highest intensity of 5.0E5 and m/z tolerance of 12 ppm. For chromatogram deconvolution, the algorithm used was the wavelets (ADAP) (Myers et al., 2017). The intensity window S/N was used as S/N estimator with a signal to noise ratio set at 10, a minimum feature height at 5.0E5, a coefficient area threshold at 130, a peak duration ranges from 0.0 to 0.5 min and the RT wavelet range from 0.01 to 0.03 min. Isotopes were detected using the isotopes peaks grouper with a m/z tolerance of 12 ppm, a RT tolerance of 0.01 min (absolute), the maximum charge set at 2 and the representative isotope used was the most intense. Each feature list was filtered before alignment to keep only features with an associated MS2 scan and a RT between 0.5 and 8.0 min using the feature filtering. At this step, individual feature tables and spectra were exported using the “export to GNPS” module to generate a MEMO matrix from unaligned samples. To generate a MEMO matrix from aligned samples, peak alignment was performed using the join aligner method (*m/z* tolerance at 40 ppm), absolute RT tolerance 0.2 min, weight for m/z at 2 and weight for RT at 1 and a weighted dot-product cosine similarity of 0.3. The aligned feature list (119,182 features) was exported using the export to GNPS module.

#### MEMO analysis

Spectra were processed using memo_ms package (0.1.3) and the following parameters for spectra import were used: min_relative_intensity: 0.01, max_relative_intensity: 1, min_peaks_required: 10, losses_from: 10, losses_to: 200, n_decimals: 2. Peaks/losses occurring in blanks samples were removed. Distance matrices were computed as described above. TMAP and UMAP were performed using tmap (1.0.0) and umap-learn (0.5.2) packages respectively after sample-wise normalization of the MEMO matrices.

#### Activity against *Trypanosoma cruzi* assay

Activity against *T. cruzi*. Rat skeletal myoblasts (L-6 cells) were seeded in 96-well microtiter plates at 2000 cells/well in 100 μL RPMI 1640 medium with 10% FBS and 2 mM l-glutamine. After 24 h the medium was removed and replaced by 100 μl per well containing 5000 trypomastigote forms of T. cruzi Tulahuen strain C2C4 containing the β-galactosidase (Lac Z) gene (Buckner et al., 1996). After 48 h the medium was removed from the wells and replaced by 100 μl fresh medium. Samples were dissolved in 5% DMSO at 0.2 mg/ml. 5 ml and 1 ml of the sample solution respectively were added to the wells. The test concentrations were 10 mg/ml and 2 mg/ml. After 96 h of incubation the plates were inspected under an inverted microscope to assure growth of the controls and sterility. Then the substrate CPRG/Nonidet (50 μl) was added to all wells. A color reaction developed within 2–6 h and could be read photometrically at 540 nm. The data were evaluated in Excel. For each test concentration, the percent growth inhibition was calculated in comparison with an untreated control. Benznidazole at 10 mg/ml was included as positive control.

#### Cytotoxicity assay: L-6 cells

Assays were performed in 96-well microtiter plates, each well containing 100 μl of RPMI 1640 medium supplemented with 1% L-glutamine (200mM) and 10% fetal bovine serum, and 4000 L-6 cells (a primary cell line derived from rat skeletal myoblasts) (Page et al., 1993; Ahmed et al., 1994). Samples were dissolved in 5% DMSO at 0.2 mg/ml. 5 ml and 1 ml of the sample solution respectively were added to the wells. The test concentrations were 10 mg/ml and 2 mg/ml. After 70 hours of incubation the plates were inspected under an inverted microscope to assure growth of the controls and sterile conditions. 10μl of Alamar Blue was then added to each well and the plates incubated for another 2 hours. Then the plates were read with a Spectramax Gemini XS microplate fluorometer (Molecular Devices Corporation, San Jose, CA, USA) using an excitation wavelength of 536 nm and an emission wavelength of 588 nm. The data were evaluated in Excel. For each test concentration, the percent growth inhibition was calculated in comparison with an untreated control. Podophyllotoxin (Sigma P4405) at 0.1 mg/ml was included as positive control.

### Additional plant extract samples

#### Description of the *Waltheria indica* extracts

These three samples correspond to the source of the waltherione derivatives, previously described in the following article for the roots (Cretton et al., 2014) and aerial parts (Cretton et al., 2016), respectively. Consult these references for further details on the extract preparation and chemical content of *Waltheria indica*. Raw and processed data have been deposited on the GNPS/MASSive repository accession number [MSV000088521](https://massive.ucsd.edu/ProteoSAFe/dataset.jsp?task=81caae4ac3724aacafaeab45ed989157).

#### LC-MS/MS analysis

*Waltheria indica* samples corresponded to additional extracts absent from the previously described 1,600 plants extract collection. For *Waltheria indica* aerial parts 2014, the initial analysis conditions are described below.

Chromatographic separation was performed on a Thermo Dionex Ultimate 3000 UHPLC system interfaced to a Q-Exactive Plus mass spectrometer (Thermo Scientific, Bremen, Germany), using a heated electrospray ionization (HESI-II) source. The LC conditions were as follows: column, Waters BEH C18 150 × 2.1 mm i.d., 1.7 μm; mobile phase, (A) water with 0.1% formic acid and (B) acetonitrile with 0.1% formic acid; flow rate, 460 μL/min; injection volume, 3 μL; gradient, linear gradient of 25%-100% B over 30 min followed by an isocratic step of 100% B for 10 min. In positive ion mode, diisooctyl phthalate C_24_H_38_O_4_ [M + H]+ ion (m/z 391.28429) was used as internal lock mass. The optimized HESI-II parameters were as follows: source voltage, 4.0 kV (pos); sheath gas flow rate (N2), 50 units; auxiliary gas flow rate, 12 units; spare gas flow rate, 2.5; capillary temperature, 266.25 °C (pos), S-Lens RF Level, 50. The mass analyzer was calibrated using a mixture of caffeine, methionine-arginine-phenylalanine-alanine-acetate (MRFA), sodium dodecyl sulfate, sodium taurocholate, and Ultramark 1621 in an acetonitrile/methanol/water solution containing 1% acid by direct injection. The data-dependent MS/MS events were performed on the 5 most intense ions detected in full scan MS (Top5 experiment). The MS/MS isolation window width was 1 *m/z*, and the stepped normalized collision energy (NCE) was set to 20–35–50 units. In data-dependent MS/MS experiments, full scans were acquired at a resolution of 35000 fwhm (at m/z 200) and MS/MS scans at 17500 fwhm both with a maximum injection time of 50 ms. After being acquired in MS/MS scan, parent ions were placed in a dynamic exclusion list for 3.0 s.

*Waltheria indica* roots and aerial parts were reprofiled in 2018 using the same analytical platform and identical LC and MS methods as used for the plant extract dataset (see corresponding Methods section).

#### Data-processing

Data were processed using MZmine as the plant extract dataset (see above), with slightly adapted parameters. For chromatogram deconvolution, the intensity window S/N was used as S/N estimator with a signal to noise ratio set at 10, a minimum feature height at 1.0E6, a coefficient area threshold at 60, a peak duration ranges from 0.0 to 0.5 min and the RT wavelet range from 0.01 to 0.03 min.

#### MEMO analysis

MEMO analysis was performed with the same parameters as the for the plant extract dataset analysis.

## Results and Discussion

### Generation of MEMO vectors and matrices

Two approaches can be followed to generate a MEMO matrix and are described in Figure 1. On one side (Figure 1 A1), it is possible to start directly from unaligned samples to construct a sample’s MEMO vector (Figure 1 B). In this case, the only required input is a .mgf file (Mascot Generic Format, a text format commonly used to report fragmentation mass spectrometry data) containing all sample’s features MS/MS spectra. These spectra can be exported from features detection software such as MZmine for example (Pluskal et al., 2010). First, MS2 spectra are filtered and binned using matchms and spec2vec packages: peaks and losses to the precursor of the spectrum are converted into words (i.e. peak@mz and loss@mz) and grouped into so-called documents (Huber et al., 2020, 2021). Documents are then aggregated by summing peaks and losses occurrences across spectra of a given sample to obtain the MEMO vector (Figure 1 B). Samples’ MEMO vectors can then be aligned to obtain a MEMO matrix (Figure 1 C). This way of building a MEMO matrix without prior RT based features alignment across samples is defined hereafter as *“MEMO from unaligned samples”* approach (Figure 1 A1).

**Figure 1:**
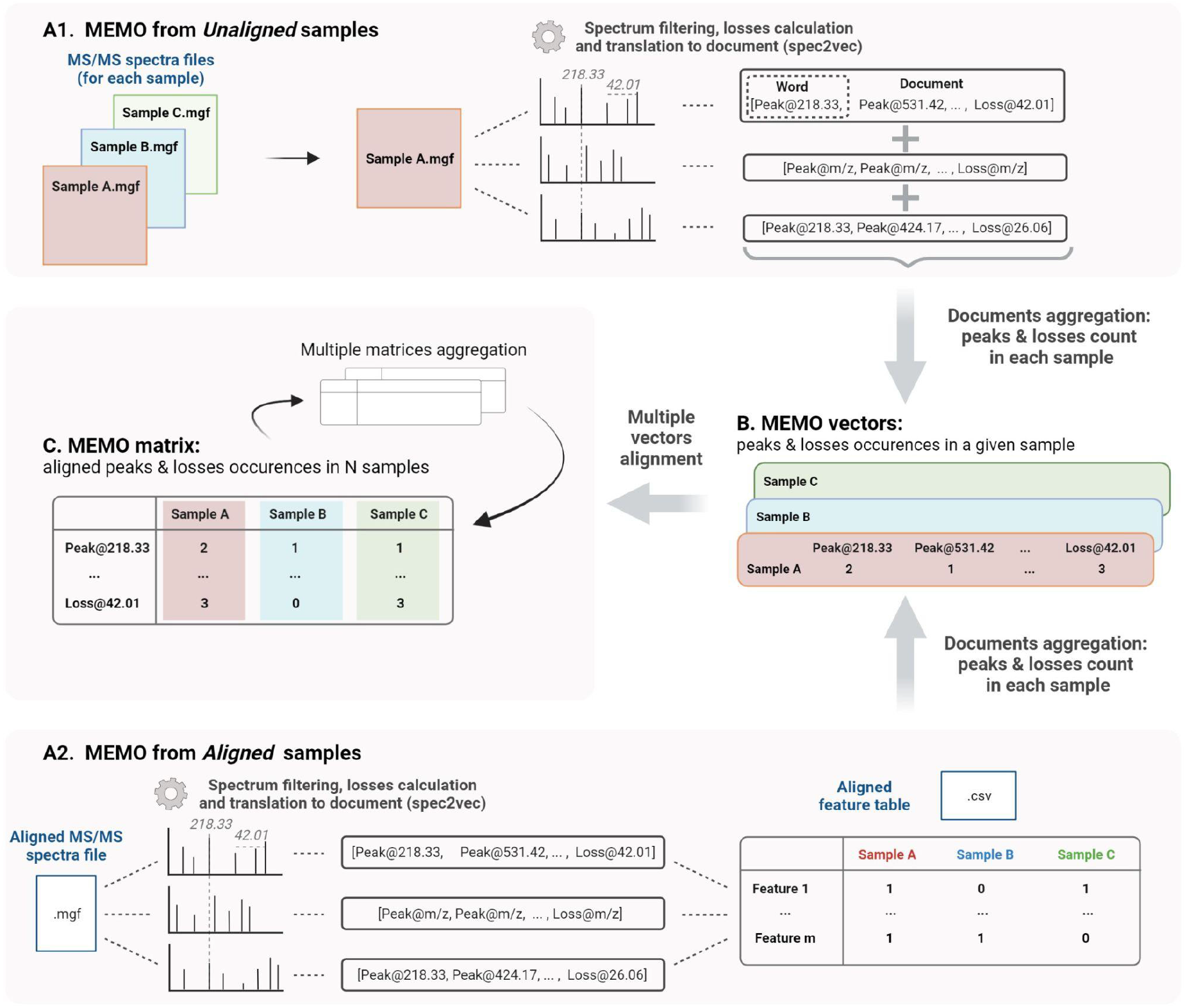
Workflow to generate a MEMO matrix from usual outputs of feature detection pipelines from unaligned samples (*MEMO from Unaligned samples*, **A1**) or from previously aligned samples (*MEMO from Aligned samples*, **A2**). For both methods, features’ MS/MS spectra are first binned and neutral losses relative to the precursor losses are calculated to build so-called *documents* using the spec2vec package (Spectrum to Document step). In (**A1)**, occurrence of each word (peak or loss) in each spectra file (i.e. sample) is counted. Words are then extracted from their Document context and aggregated to build a MEMO vector for each sample (**B**). In (**A2**), the building of the MEMO vector (**B**) is done using the Aligned Feature Table to determine whether a feature is detected or not in each sample. Likewise, each word is counted within the originating sample and extracted from its Document (or spectral) context. MEMO vectors are then aligned on words in a so-called MEMO matrix (**C**). These MEMO matrices can then be (optionally) aggregated. Figure created using biorender.com.

On the other side (Figure 1 A2), it is also possible to start the process from an aligned feature intensity table (.csv file) and corresponding MS/MS spectra file (.mgf file). These files are usually generated by metabolomics workflows such as the Feature-Based Molecular Networking analysis (FBMN) (Nothias et al., 2020). In this case, the occurrence of each peak and loss within a given sample is counted using the feature intensity table to determine whether a feature is detected or not into the sample (Figure 1 A2). Note here that it is not the intensity table of the MS1 feature (peak intensity or area) that is considered but rather a simple presence/absence matrix. This *“MEMO from aligned samples”* workflow can conveniently complement pre-existing metabolomics data treatment pipelines (Figure 1A2).

The main advantage of using unaligned files to construct a MEMO matrix is that the time-consuming feature alignment step, which is prone to batch effects, can be completely bypassed. This approach is thus well suited to the analysis of large data sets and is, to the best of our knowledge, the only way to compare samples acquired with different LC methods. Multiple MEMO vectors and/or MEMO matrices can be merged, facilitating the subsequent addition of supplementary samples’ MEMO vectors. This is a convenient characteristic of the MEMO analytical workflow, which allows iterative analysis of evolving datasets (see section 4, Application of MEMO on a large plant extracts dataset: batch effect correction and drug-discovery application).

### Evaluation of the MEMO method

#### Comparison to state-of-the-art MS/MS-based clustering methods

To benchmark the MEMO approach, we compared it to the two previously established MS/MS based sample clustering methods: CSCS and Qemistree (Sedio et al., 2017; Tripathi et al., 2021). CSCS combines the cosine pairwise similarity between spectra with the feature table to compute a distance matrix among samples while Qemistree relies on predicted molecular fingerprints (via SIRIUS/CSI:FingerID) to generate a tree used for UniFrac distance computation among samples (Duhrkop et al., 2015, 2019). To perform the comparison of the different methods, we used an experimental dataset previously built for the benchmarking of the Qemistree tool. This dataset is constituted by 27 samples, which were acquired in triplicates using two different LC methods (named C18 and C18 RT-shift) thus simulating a RT shift and a strong artificial batch effect. It was also acquired on two different mass spectrometers (Q-exactive Orbitrap (QE) and Maxis II q-TOF (q-ToF)). The composition of the samples was the following: four chemodiverse “parent samples” (one plasma, one tomato and two feces samples), binary mixtures (in different proportions) and quaternary mixtures of these four “parent samples”. For more details, see (Tripathi et al., 2021). Hereafter, we refer to this dataset as the “evaluation dataset”.

The objective of this comparison was to evaluate each clustering method’s ability to mitigate retention time shift effect. For this, we applied six different techniques - *MEMO from unaligned* and *aligned samples*, *Qemistree weighted/unweighted* and *CSCS weighted/unweighted* - to the samples analyzed with the two different LC-methods on the Q-Exactive MS. We also added the results of an MS/MS agnostic clustering method (*Feature Table Bray-Curtis*) solely based on the MS1 feature table intensity to serve a baseline. The overall objective of these clustering methods is to capture a maximum of the compositional (dis)similarity across extracts despite the strong experimental batch-effect induced by the artificial RT-shift. Results are presented in Figure 2.

**Figure 2:**
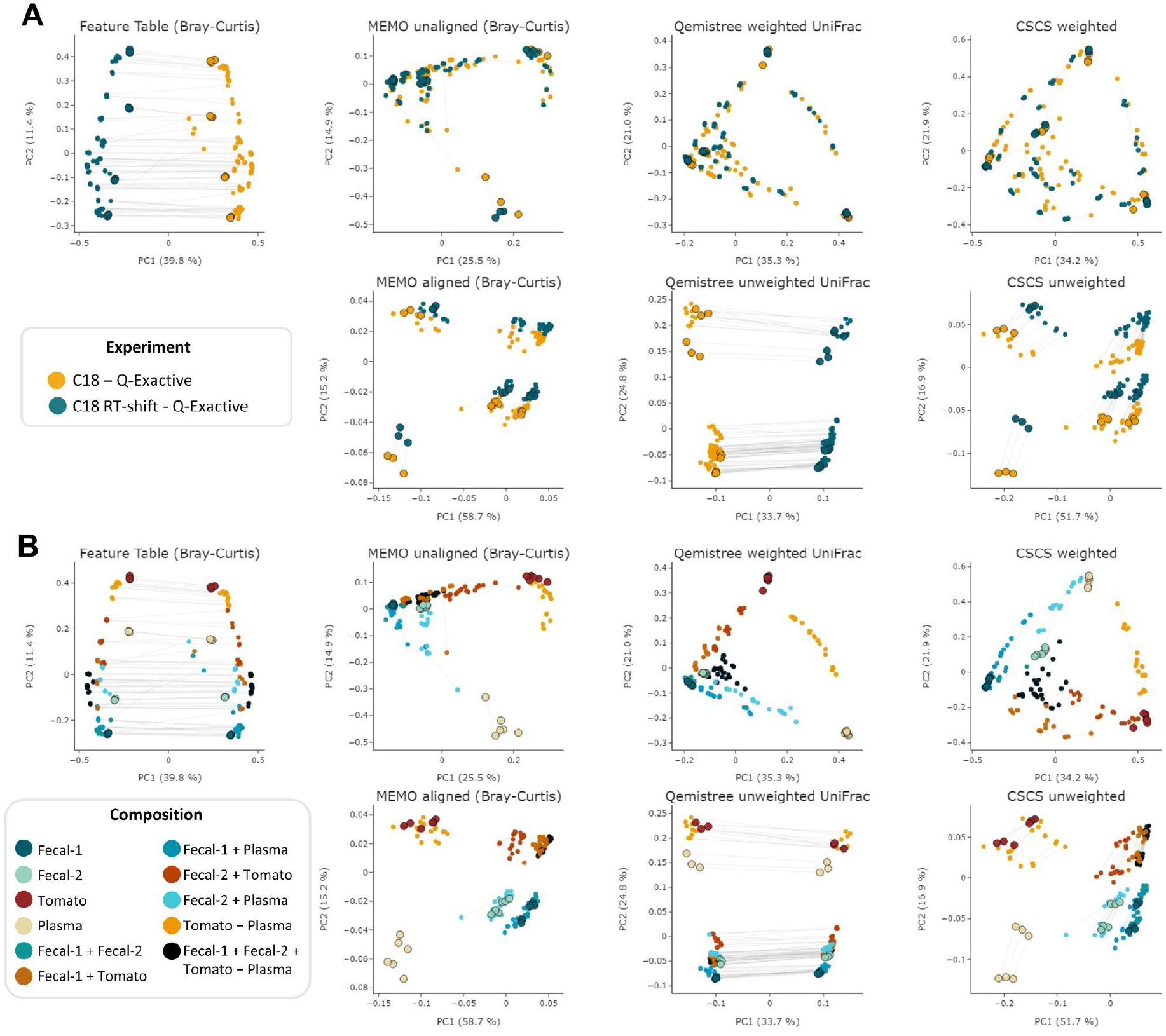
PCoA comparison of a classical and MS/MS agnostic approach (*Feature Table Bray Curtis*) and three MSMS informed clustering approaches (the *MEMO from unaligned/aligned* (Bray-Curtis distance), *Qemistree weighted/unweighted* (UniFrac distance) and *weighted/unweighted CSCS*) on the evaluation dataset acquired using 2 different LC methods (C18 and C18 RT-shift) on the Q-Exactive mass spectrometer. Samples are colored according to the used experimental LC method (**A**) or their composition (**B**). For statistical analysis, see Table 1. The samples corresponding to the same mixture and replicate in each dataset (C18 and C18 RT-shift) are linked (gray line). Parent samples are bigger and have a black border line. Interactive visualizations are available [here](https://mandelbrot-project.github.io/memo_publication_examples/).

The observation of these results first indicates the importance of using fragmentation spectra in order to decrease the RT shift effect. Indeed, methods considering mass fragmentation information, with the exception of Qemistree unweighted, all appear to mitigate the artificial batch effect (Figure 2 A).

Secondly, it is to be noticed that, overall, similar clustering patterns are observed between *MEMO from unaligned samples, Qemistree weighted* and *CSCS weighted*, on one side, and *MEMO from aligned samples* and *CSCS unweighted* on the other side. For *MEMO from unaligned, Qemistree weighted* and *CSCS weighted*, mixtures of the same parent samples but of different proportions can be distinguished, and the best performing method is *Qemistree* (Figure S1 and S2). Interestingly, *MEMO from aligned samples* is closer to *CSCS unweighted* than to *MEMO from unaligned samples*. This could be explained by the fact that a gap-filling was applied after alignment, causing a decrease of the impact of the proportions of the parent samples in the mixtures when compared to *MEMO from unaligned samples*.

To confirm the previous visual interpretations, a statistical analysis was performed using the permutational analysis of variance (PERMANOVA) test (Anderson, 2017) (See Table 1). Overall, larger pseudo-F values indicate more pronounced group separation. The optimal clustering approach in our example case should thus ideally have the smallest pseudo-F value when considering a grouping via the *experiment* factor (the RT induced batch effect) and the larger pseudo-F value when considering the *composition* factor. First, the observed pseudo-F value when considering the *experiment* parameter (C18 vs C18 RT-shift) is clearly smaller for MS2 based methods (pseudo-F <= 81.64) compared to the classical intensity-based dissimilarity analysis (pseudo-F = 113.55). This confirms, as expected, a much stronger influence of this experiment parameter when features are considered as independent variables without MS/MS inclusion. The opposite effect is observed when measuring the pseudo-F value on the *composition* parameter: values are higher in MS2 based methods (>=11.10) than in the *feature table Bray-Curtis* analysis (4.72), confirming the visualization of the first two Principal Coordinates in Fig. 1. The difference between *MEMO from unaligned* and *aligned samples* is important, highlighting the important influence of the alignment and gap-filling steps. While *MEMO from unaligned samples* appears to be the most efficient at mitigating batch effect, its ability to discriminate samples according to their content is lower than *MEMO from aligned samples*, at least statistically. It is however to be noticed that since binary mixtures are constituted of different proportions of the parent samples, mixtures groups are heterogeneous and this impacts the results of the PERMANOVA test.

**Table 1:** PERMANOVA results for the different evaluated metrics on two categorical attributes; the experiment (2 groups: C18, C18-RTshift) and the composition (11 groups: Fecal-1, Fecal-2, Tomato, Plasma, Fecal-1 + Fecal-1, Fecal-1 + Tomato, Fecal-1 + Plasma, Fecal-2 + Tomato, Fecal-2 + Plasma, Tomato + Plasma, Fecal-1 + Fecal-2 + Tomato + Plasma). The optimal clustering approach should ideally have the smallest pseudo-F value when considering a grouping via the *experiment* factor (the RT induced batch effect) and the larger pseudo-F value when considering the *composition* factor.

| Metric | Group: Experiment |  | Group: Composition |  |
| --- | --- | --- | --- | --- |
|  | pseudo-F | p-value | pseudo-F | p-value |
| Feature table (Bray-Curtis) | **113.55** | 0.001 | **4.72** | 0.001 |
| MEMO from unaligned samples (Bray-Curtis) | **2.23** | 0.017 | **15.51** | 0.001 |
| MEMO from aligned samples (Bray-Curtis) | **25.39** | 0.001 | **62.79** | 0.001 |
| Qemistree weighted UniFrac | **18.22** | 0.001 | **49.54** | 0.001 |
| Qemistree unweighted UniFrac | **81.64** | 0.001 | **11.10** | 0.001 |
| CSCS weighted | **17.94** | 0.001 | **52.53** | 0.001 |
| CSCS unweighted | **28.27** | 0.001 | **62.10** | 0.001 |

A strong advantage of the MEMO approach is that it is fast and computationally cheap compared to the two other metrics (Table 2). A strict comparison of the computation time of these three conceptually different and diversely implemented methods is complicated. However, a comparison of the methods from the users’ practical point of view is doable and of interest. On large datasets, such as the plant extract dataset (1920 samples and 906,509 MS/MS spectra), a MEMO matrix can be computed rapidly (less than 10 minutes) on a simple laptop (see specifications in Table 2 caption). In comparison, CSCS required much more computational time (295 min) despite being parallelized on a more powerful machine (see specifications in Table 2 caption). Computational requirements for the Qemistree approach are high and its application on such a dataset is complicated. The efficiency of the MEMO approach offers exciting perspectives for repository-scale analyses of large spectral ensembles.

**Table 2:** Computational time required to obtain the distance matrix used for PCoA analysis for each of the compared metric. The feature detection step (using MZmine 2) is not considered since it is the same for all three methods. The time for uploading, downloading and unzipping the data when needed is not taken into account. All 7032 features were considered for MEMO and CSCS analyses. *Windows 10 64 bits, Intel Core i7-8750H, 2.20 GHz, 6 cores, 16 Gb of RAM - no parallelization. ** Ubuntu 20.04 LTS 64 bits, AMD Ryzen Threadripper 3970X (3.70 GHz / 128 MB), 32 cores, 256 Gb of RAM - parallelized on 40 threads

| Dataset | Metric | GNPS<br>FBMN<br>(min) | GNPS<br>Qemistree<br>(min) | Local<br>computation<br>(min) | Total<br>(min) |
| --- | --- | --- | --- | --- | --- |
| Evaluation<br>dataset (198<br>samples) | MEMO from<br>*unaligned* samples | NA | NA | 1.1* | **1.1** |
|  | MEMO from<br>*aligned* samples | NA | NA | 0.5* | **0.5** |
|  | CSCS weighted | 49 | NA | 13.3** | **62.3** |
|  | CSCS unweighted | 49 | NA | 13.3** | **62.3** |
|  | Qemistree<br>weighted | 49 | 796 | < 0.1* | **845.0** |
|  | Qemistree<br>unweighted | 49 | 796 | < 0.1* | **845.0** |
| Plant extracts<br>dataset (1,920<br>samples) | MEMO from<br>unaligned samples | NA | NA | 9.9* | **9.9** |
|  | CSCS weighted | 45 | NA | 250.4** | **295.4** |

#### Organizational capability for unaligned samples analyzed on two different mass spectrometers

To evaluate MEMO’s ability to cluster similar samples affected by an alternative batch effect, explained this time by different MS platforms, we applied it to the evaluation dataset samples analyzed on a Q-Exactive Orbitrap (QE) and a Quadrupole-Time of Flight (Q-ToF) MS. A MEMO matrix was generated from aligned samples for each platform, and the two resulting matrices were then aligned. The counts in the final MEMO matrix were normalized sample-wise to mitigate effects of peaks/loss occurrence difference between MS. The PCoA performed on the final matrix is presented in Figure 3.

**Figure 3:**
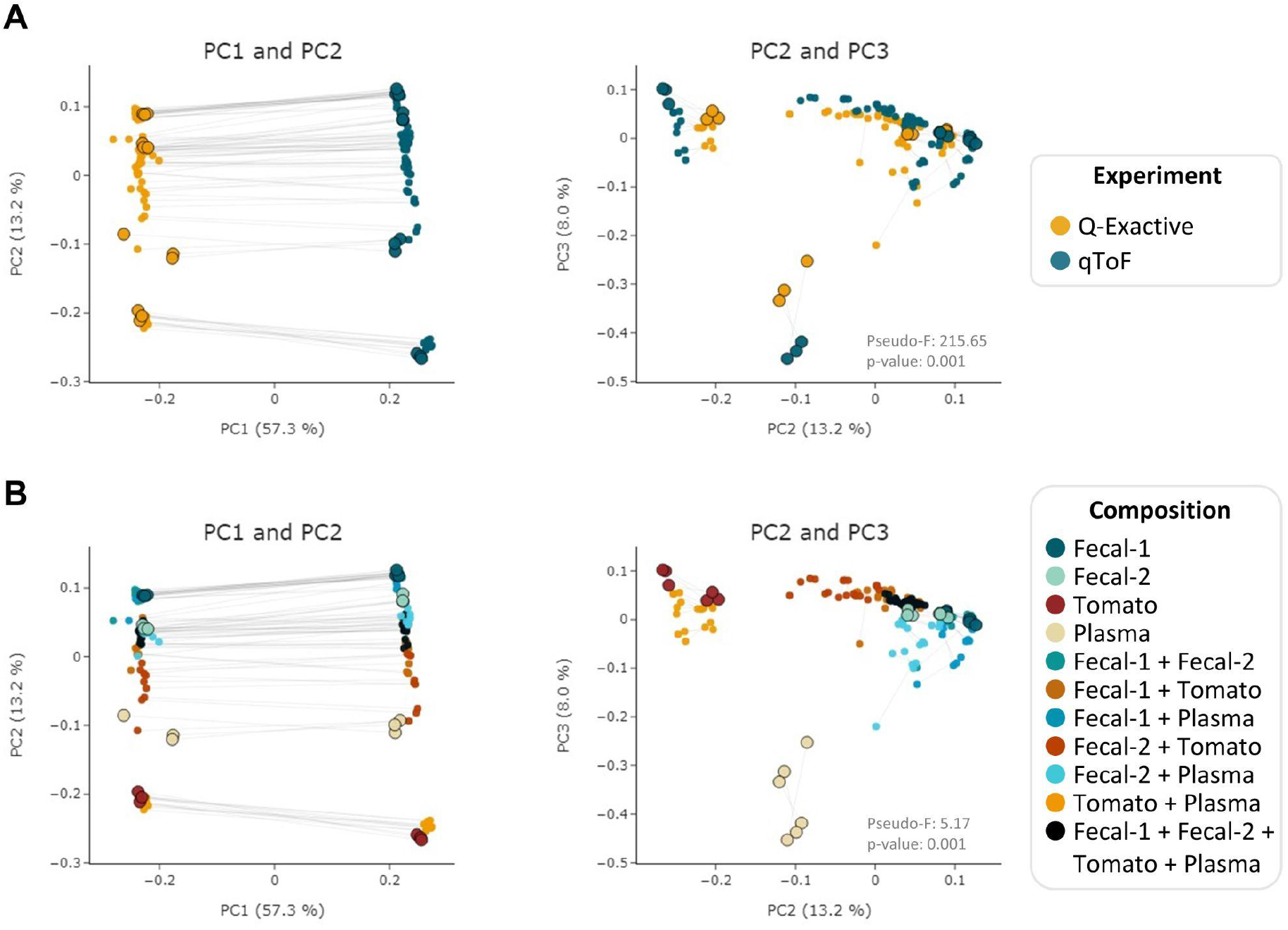
PCoA (Bray-Curtis) on the MEMO matrix analysis of the Qemistree samples analyzed on a Q-ToF and an Q-Exactive orbitrap MS. The samples corresponding to the same mixture and replicate in each dataset are linked (gray line) and parent samples are bigger and have a black border line. Samples are colored according to the used *experiment* (**A**) or their *composition* (**B**). PERMANOVA: Experiment: pseudo F= 215.65, P-value = 0.001; Composition: pseudo F= 5.17, P-value = 0.001. Interactive visualizations are available [here](https://mandelbrot-project.github.io/memo_publication_examples/).

When looking at the first two PCs, the strongest factor driving the clustering is, as expected, related to the MS platform. This is confirmed by the PERMANOVA results with a pseudo-F value of 213.35 (p-value = 0.001) for the *experiment* factor (Figure 3 A). However, a good clustering according to samples’ composition can be observed on PC2 and PC3 (pseudo-F = 5.19, p-value = 0.001 for the *composition* factor) (Figure 3B). The possibility to organize such datasets - even if imperfectly - is of interest since, to the best of our knowledge, methods to compare samples acquired on 2 different MS technologies has not been reported to date. The strong batch effect observed is certainly caused by the difference in fragmentation technique between the two instruments (higher-energy C-trap dissociation in QE vs collision-induced dissociation in Q-ToF). However, in the era of open data, methods that allow such comparison are of high value to extract knowledge from MS metabolomics studies available on public repositories such as MASSive (Wang et al., 2016) or Metabolights (Haug et al., 2020). Such comparison is possible using the ReDU framework (Jarmusch et al., 2020) but exploits the presence of annotated compounds within each sample. MEMO, being annotation-independent, is thus a complementary holistic view of the data considered.

### Application of the MEMO method to a large plant extract dataset: batch effect correction and implementation in drug discovery

To confirm MEMO’s scalability, we applied it to a large plant extract dataset (1,600 extracts from 767 botanical species and different plant parts) analyzed in our lab over a year span in 19 batches. The resulting dataset is public and can be accessed through the GNPS/MASSive repository MSV000087728. First, as depicted on Figure 4, we could observe that this dataset suffers from a strong batch effect related to the date of analysis. This batch effect is clearly visible on the Feature Table based PCoA with on one side samples analyzed before 2017-10-03, and on the other side samples analyzed after 2017-10-26 (Figure 4 A and Figure S4). This batch effect is due to both a RT shift and a MS sensitivity change, as it can be observed on 2 representative quality control (QC) samples (Figure S3). When performing the PCoA on MEMO from aligned and unaligned matrices, the influence of this batch effect is, in both cases, clearly mitigated (Figure 4 A). It is noteworthy that an injection date effect is still visible on the UMAP and TMAP visualizations, highlighting the difficulty to eliminate batch related variation (Figure S4). Since the *MEMO from unaligned sample* workflow is faster to compute than *MEMO from aligned sample* (by bypassing the feature alignment step), it will be used for the rest of the work.

**Figure 4:**
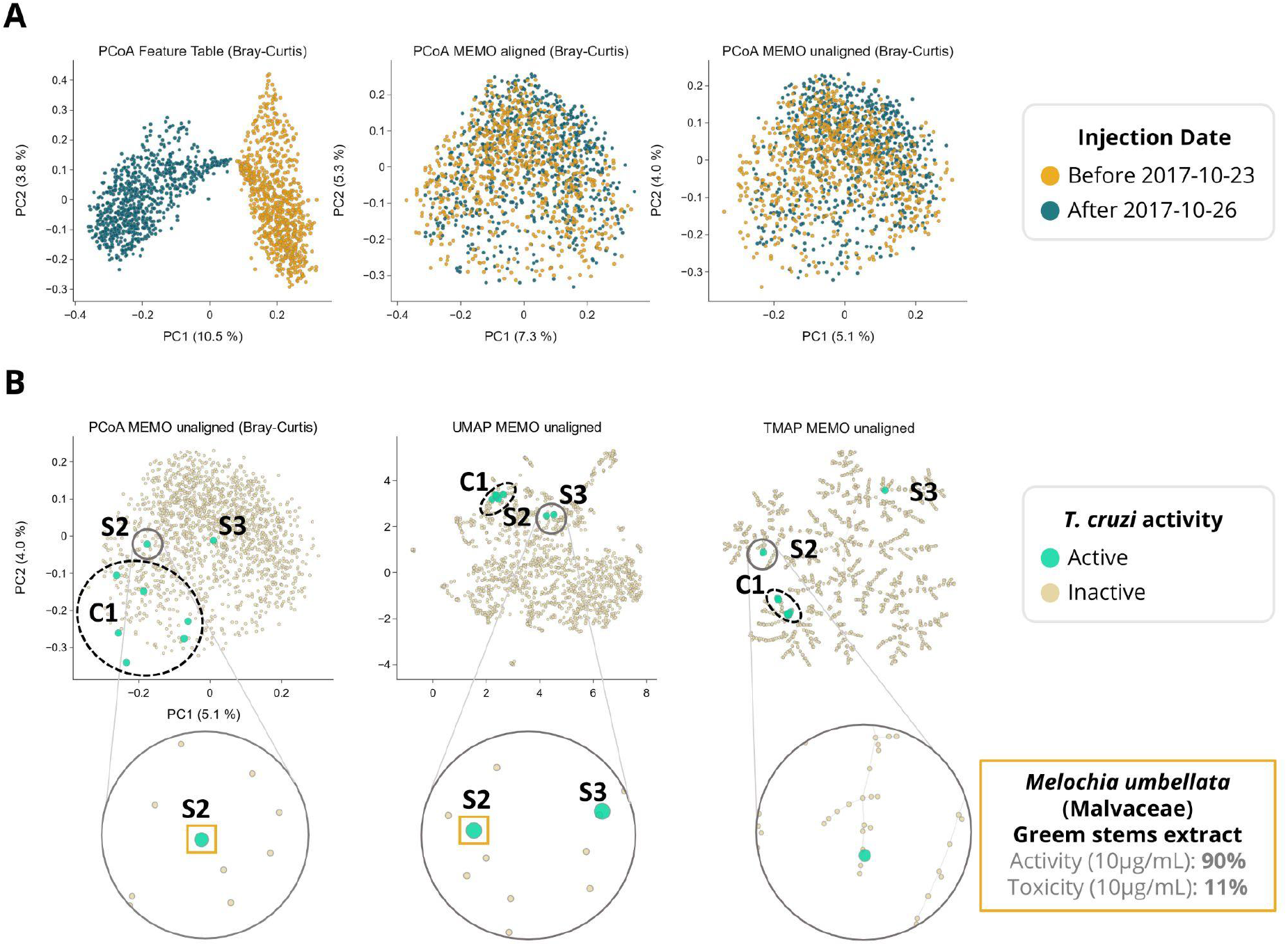
Classical MS1 intensity feature table based clustering and MEMO-based organization of a large chemo-diverse plant extracts dataset. Due to practical constraints, the 1,600 samples of the dataset were acquired over 16 months and a strong batch effect could be observed when using classical PCoA visualization on the feature table (**A**, PCoA Feature Table). This batch effect is much lower when PCoA is performed both from aligned and unaligned MEMO matrices (**A**, PCoA *MEMO from aligned* and *unaligned*). In (**B)**, the anti-*Trypanosoma cruzi* activity of the extracts is mapped on three different visualizations of the unaligned MEMO matrix (PCoA, UMAP and TMAP). An extract was classified as active if inhibition growth inhibition of *T. cruzi* was above 80% of the control and cytotoxicity against L6 cells was below 50% of the control. This comparison highlights the necessity of complementary dimensionality reduction techniques in order to gain chemical insights about the sample. Indeed, using UMAP and TMAP, one cluster (C1) of six active samples and 2 singletons samples (S2 and S3) can be observed on both the UMAP and the TMAP but not on the PCoA. C1 and S3 will be investigated in future works while S2, corresponding to *Melochia umbellata* green stems extract, has been investigated and detailed in the current work (Figure 5). Interactive visualizations are available [here](https://mandelbrot-project.github.io/memo_publication_examples/).

MEMO also enabled a good clustering of the samples according to their metabolic profile. Indeed, in the frame of a drug discovery project, we evaluated the activity of these 1,600 plant extracts against *Trypanosoma cruzi* (*T. cruzi*). In this screening, eight extracts displayed an interesting activity against *T. cruzi* (> 80% of parasite growth inhibition) coupled to a moderate cytotoxicity (< 50%). Among these eight bioactive extracts, six are clustered together (C1) and two appeared as singletons (S1 and S2) in the UMAP and TMAP visualization (Figure 4 B and Figure S5). Getting similar clustering using two different algorithms strengthens the hypothesis that observed clusters have, in addition to a biological relevance, a chemical relevance and are not visualization artifacts. This hypothesis is strengthened for C1 by the fact that among the eight extracts, six belong to the Leguminosae botanical family (three different species) and that a rapid dereplication allowed identification of structurally similar compounds in all of them. The isolation of the active compounds of the six active samples (C1) is the object of an ongoing work and will be described in a future article.

Among the two active samples not clustered in C1, *Melochia umbellata* (Houtt.) Stapf (Malvaceae) green stems extract (S2, Figure 4 B) displayed an interesting anti-trypanosomatid activity (90% growth inhibition at 10 μg/mL). Interestingly, colleagues from the University of Geneva previously discovered potent trypanocidal waltherione derivatives from another Malvaceae, *Waltheria indica* L. (Cretton et al., 2014, 2015, 2020). To evaluate whether the activity of these two plants of the same botanical family could be explained by structurally similar compounds, we added the MEMO vectors of three additional *Waltheria indica* extracts LC-MS/MS profiles, to the global MEMO matrix of the 1,600 plant extract dataset. These three *Waltheria indica* extracts profiles were heterogeneous: a first *Waltheria indica* aerial parts extract was profiled in 2014 and two *Waltheria indica* roots and aerial parts were reprofiled in 2018. Since MEMO vector based organization can be performed without the requirement for a prior alignment step, the addition of samples to a pre-existing dataset is straightforward. In this case, this was done in minutes after the processing of the individual *Waltheria indica* metabolite profiles using MZmine. Moreover, if the two 2018 samples were analyzed on the same platform with the same methods (LC and MS/MS) as the ones used for the profiling of the plant extract dataset, the 2014 sample was acquired using a different LC-gradient (40 min run on a 2.1 × 150 mm column versus 8 min run on 2.1 × 50 mm column) and a different MS (QE-Plus vs. QE-Focus orbitrap). Despite these differences, both UMAP and TMAP visualizations present a clear clustering of the three additional *Waltheria indica* samples with the trypanocidal *Melochia umbellata* green stems extract of the global plant extract dataset (Figure 5 A and B). In depth exploration of the *Melochia umbellata* green stems extract MS/MS spectra led to the identification of waltherione G, a potent anti-*Trypanosoma cruzi* compound previously isolated from *Waltheria indica* and present in the three injected extract (Figure 5 C). This identification was possible by comparing MS/MS spectra and retention time to an isolated standard of waltherione G (GNPS Library ID CCMSLIB00006718062). Despite the fact that waltherione G has not been isolated from *Melochia umbellata* until now, several related quinolone analogues have been previously reported in the genus (see Wikidata query https://w.wiki/4bWK, (Rutz et al., 2021)), further supporting this observation.

**Figure 5:**
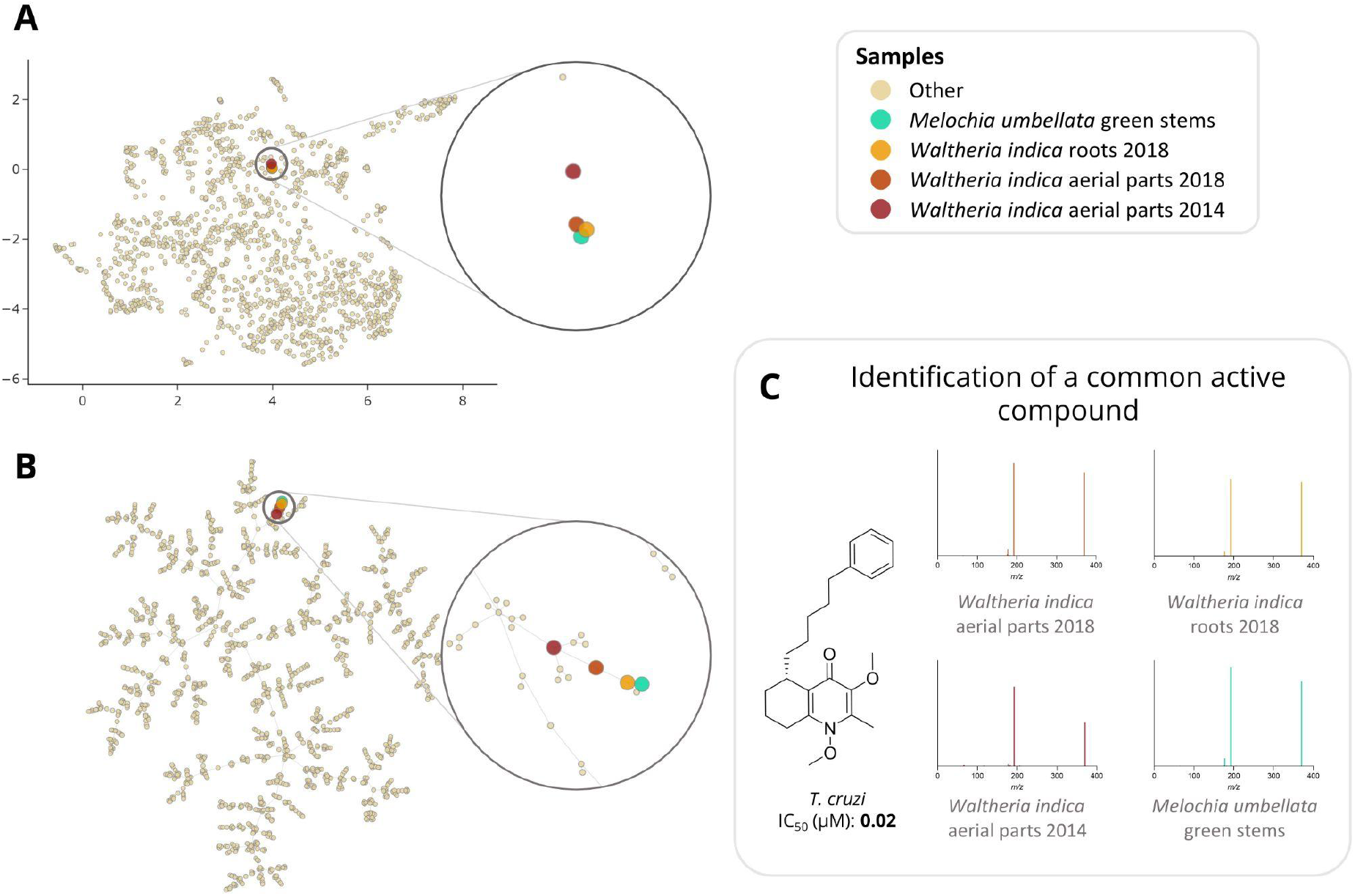
UMAP (**A**) and TMAP (**B**) visualization of the 1600 plant extracts dataset with the addition of 3 extracts rich in waltherione derivatives from *Waltheria indica* (roots 2018, aerial parts 2018 and aerial parts 2014). In (**C)**, an example of a common potent trypanocidal compound (IC_50_ = 0.02 μM), waltherione G, isolated from *Waltheria indica* and identified in all 4 extracts is shown with the corresponding MS/MS spectra from each sample. Identification of waltherione G in *Melochia umbellata* (level 1 identification of the Metabolomics Standards Initiative (Sumner et al., 2007)) was confirmed by retention time comparison with a standard (Cretton et al., 2014). Mirror view of the 2 spectra (waltherione G isolated standard and waltherione G detected in *Melochia umbellata*) is available on the Metabolomics Spectrum Resolver Web Service here (Bittremieux et al., 2020). Interactive visualizations are available [here](https://mandelbrot-project.github.io/memo_publication_examples/).

This last example illustrates how the MEMO approach can be efficiently employed to compare samples acquired in different batches, but also on different analytical platforms through quick alignment of additional MEMO vectors to a pre-established MEMO matrix. The rapid identification of the compounds likely responsible for the activity against *T. cruzi* of the *Melochia umbellata* green stems extract exemplifies how this strategy can be used for the efficient study of large natural extract collection in the frame of drug discovery projects. In further developments, techniques relying on motifs in mass spectra, such as MS2LDA, could be adapted and used to retrieve common patterns in clustered samples (van der Hooft et al., 2016). Such an implementation could facilitate the extraction of features of interest among samples. Furthermore, the MEMO vector structure allows the conversion of complex and multidimensional datasets of untargeted mass spectrometry acquired on large and chemodiverse collections of extracts to a fingerprint format perfectly amenable for future machine-learning based investigations - for example to explore the links between bioactivities of extracts and corresponding MEMO vectors.

## Conclusion

Methods exploiting fragmentation spectra to cluster samples from LC-MS/MS experiments lower batch effect and enable a better clustering than classical approaches based on aligned feature tables. However, due to technical constraints, existing strategies do not scale up easily. Here, we describe a new **M**S2 Bas**E**d Sa**M**ple Vect**O**rization (MEMO) approach, which captures the spectral diversity of complex samples in a retention time-agnostic fashion and thus allows a fast and efficient comparison of large amounts of samples without the need of a preliminary feature alignment step. The MEMO method was comparable to current state-of-the-art techniques in terms of sample clustering but with far less computational requirements. As an application example, MEMO was run on a large and chemodiverse plant extract dataset profiled over the span of a year. In less than 10 minutes and a classical laptop, MEMO was able to organize over 1,600 extracts while mitigating a strong batch effect. The exploitation of the MEMO matrix allowed to rapidly identify a structural family of bioactive compounds via the posterior aggregation to the dataset of well-studied plant extracts profiled on different chromatographic and mass spectrometric platforms. MEMO significantly advances the field of large-scale sample comparison and should facilitate knowledge extraction from the ever-increasing corpus of data generated by the metabolomics community nowadays.

## Data availability statement

MEMO is available as a python package and can be found on https://github.com/mandelbrot-project/memo. It can also be installed through https://pypi.org/project/memo-ms/. For the evaluation dataset, data and methods have been deposited on the GNPS/MASSive repository under accession number [MSV000083306](https://massive.ucsd.edu/ProteoSAFe/dataset.jsp?task=c5312f8ecb7d45feb35a723ba68874d2). The GNPS Feature-Based Molecular Networking (FBMN) job is available [here](https://gnps.ucsd.edu/ProteoSAFe/status.jsp?task=044e981ff0d84246ae5c91ef3db643a8) and the GNPS Qemistree job is available [here](https://gnps.ucsd.edu/ProteoSAFe/status.jsp?task=8ca56d6e33bc4106b46ba5e3510c91cb). For the plant extract dataset, raw and processed data have been deposited on the GNPS/MASSive repository accession number MSV000087728. The samples’ metadata are also available on the MASSive repository [here](https://massive.ucsd.edu/ProteoSAFe/dataset_files.jsp?task=b753bf1e39cb4875bdf3b786e747bc15#%7B%22table_sort_history%22%3A%22main.collection_dsc%22%2C%22main.attachment_input%22%3A%22updates%2F2021-12-21_pmallard_cde57c76%22%2C%22main.collection_input%22%3A%22metadata%7C%7CEXACT%22%7D). For the *Waltheria indica* samples, raw and processed data have been deposited on the GNPS/MASSive repository accession number [MSV000088521](https://massive.ucsd.edu/ProteoSAFe/dataset.jsp?task=81caae4ac3724aacafaeab45ed989157). All scripts used for data analysis are available on [Github](https://github.com/mandelbrot-project/memo_publication_examples).

## Acknowledgments

The authors are grateful to Green Mission Pierre Fabre, Pierre Fabre Research Institute, Toulouse, France and especially to Bruno David for establishing and sharing this unique library of extracts. The authors thank the Drugs for Neglected Diseases initiative for having financially supported the biological evaluation of the Pierre Fabre collection in *T. cruzi* and L-6 cellular assays at Swiss Tropical and Public Health Institute. JLW, LFN and PMA are thankful to the Swiss National Science Foundation for the funding of the project (SNF N° CRSII5_189921 / 1). The authors are grateful to the authors of the Qemistree article for making their valuable evaluation dataset available to the community, allowing for optimal data reuse.

## Author contributions

AG and PMA conceptualized the method and designed the study. AG, SC, LFN and PMA acquired and gathered the data. MK performed the biological assays. AG and PMA performed the data analysis and visualization. AG and PMA wrote the original manuscript. AG, PMA and JLW revised the manuscript. AG, FH and PMA developed the memo_ms python package. All authors read, reviewed and approved the article.

## Supplementary Material

**Figure S1:**
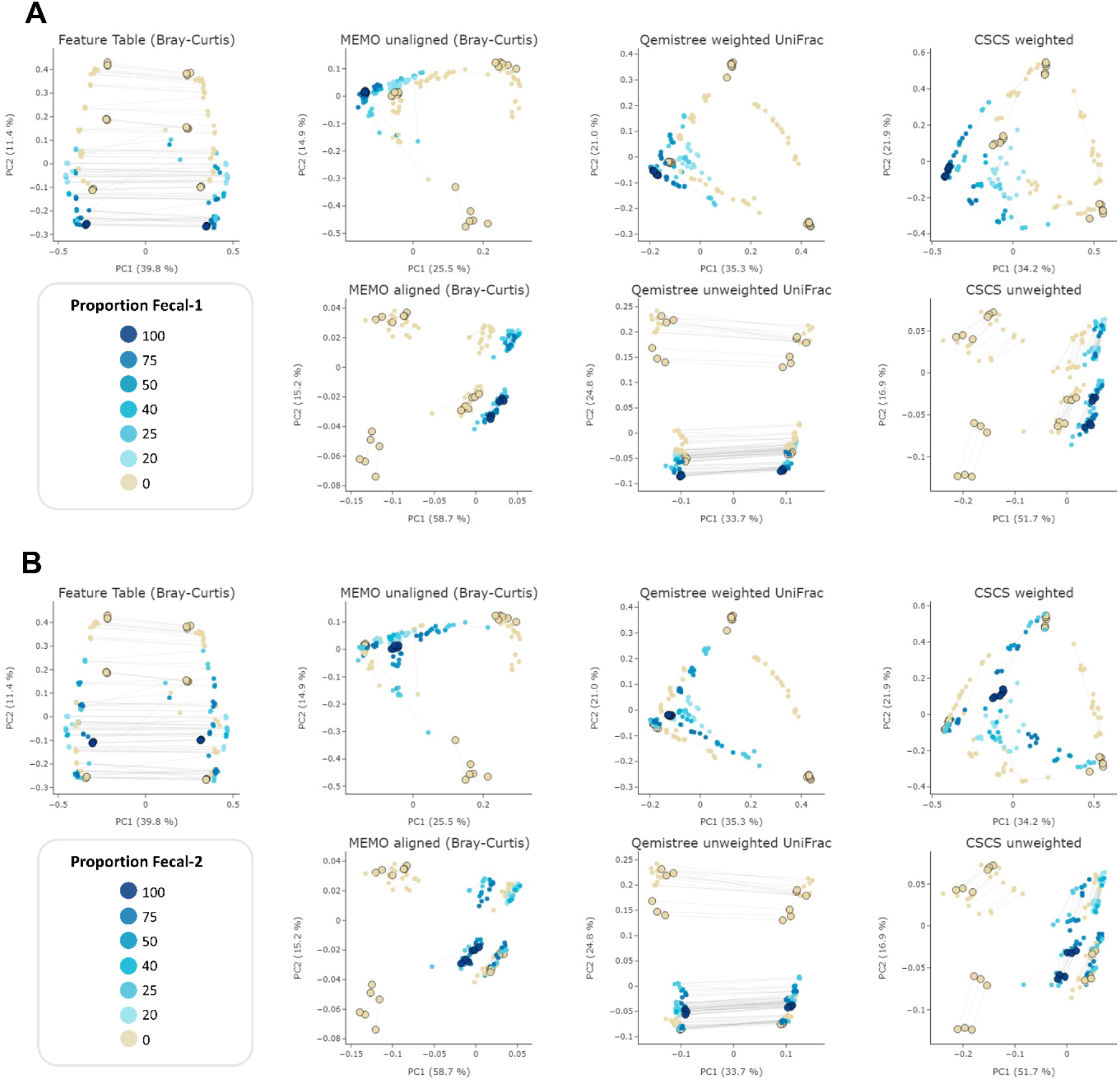
PCoA comparison of a classical and MS/MS agnostic approach (Feature Table Bray Curtis) and three MSMS informed clustering approaches (the MEMO from unaligned/aligned (Bray-Curtis distance), Qemistree weighted/unweighted (UniFrac distance), weighted/unweighted CSCS and Feature-Table (Bray-Curtis distance) clustering) on the evaluation dataset acquired using 2 different LC methods on the Q-Exactive mass spectrometer (C18 and C18 RT-shift). Samples are colored according to the proportion of fecal-1 sample (**A**) or fecal-1 sample (**B**). For statistical analysis, see Table 1. The samples corresponding to the same mixture and replicate in each dataset (C18 and C18 RT-shift) are linked (gray line). Parent samples are bigger and have a black border line. Interactive visualizations are available [here](https://mandelbrot-project.github.io/memo_publication_examples/).

**Figure S2:**
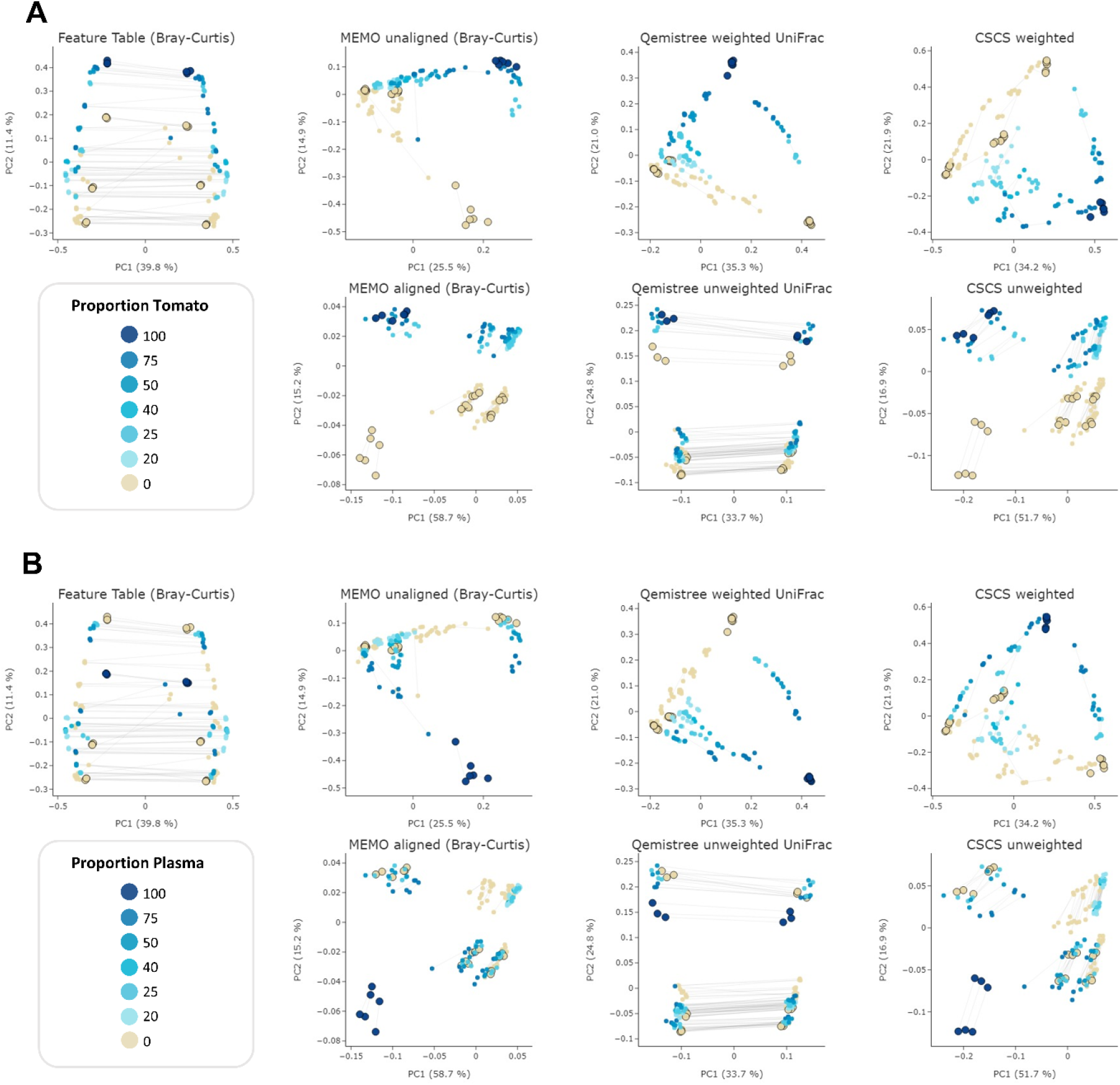
PCoA comparison of a classical and MS/MS agnostic approach (Feature Table Bray Curtis) and three MSMS informed clustering approaches (the MEMO from unaligned/aligned (Bray-Curtis distance), Qemistree weighted/unweighted (UniFrac distance), weighted/unweighted CSCS and Feature-Table (Bray-Curtis distance) clustering) on the evaluation dataset acquired using 2 different LC methods on the Q-Exactive mass spectrometer (C18 and C18 RT-shift). Samples are colored according to the proportion of tomato sample (**A**) or plasma sample (**B**). For statistical analysis, see Table 1. The samples corresponding to the same mixture and replicate in each dataset (C18 and C18 RT-shift) are linked (gray line). Parent samples are bigger and have a black border line. Interactive visualizations are available [here](https://mandelbrot-project.github.io/memo_publication_examples/).

**Figure S3:**
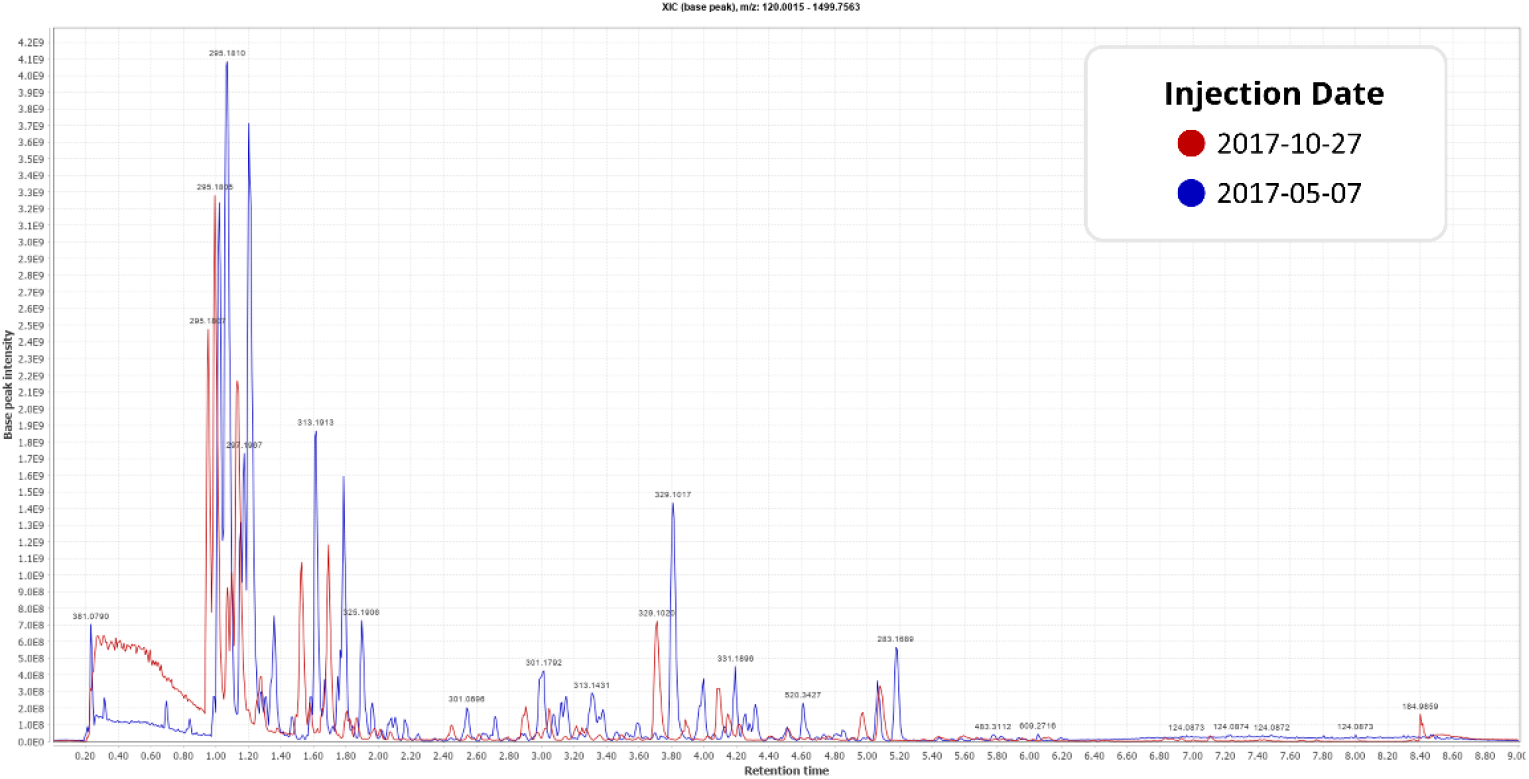
TIC ESI-(+) of two representative QC samples from each of the two observed batches. QC sample is a mixture of 5 plant ethyl acetate extracts (*Arnica montana*, *Cinchona succirubra* (syn. *Cinchona pubescens*), *Ginkgo biloba*, *Panax ginseng*, *Salvia officinalis*).

**Figure S4:**
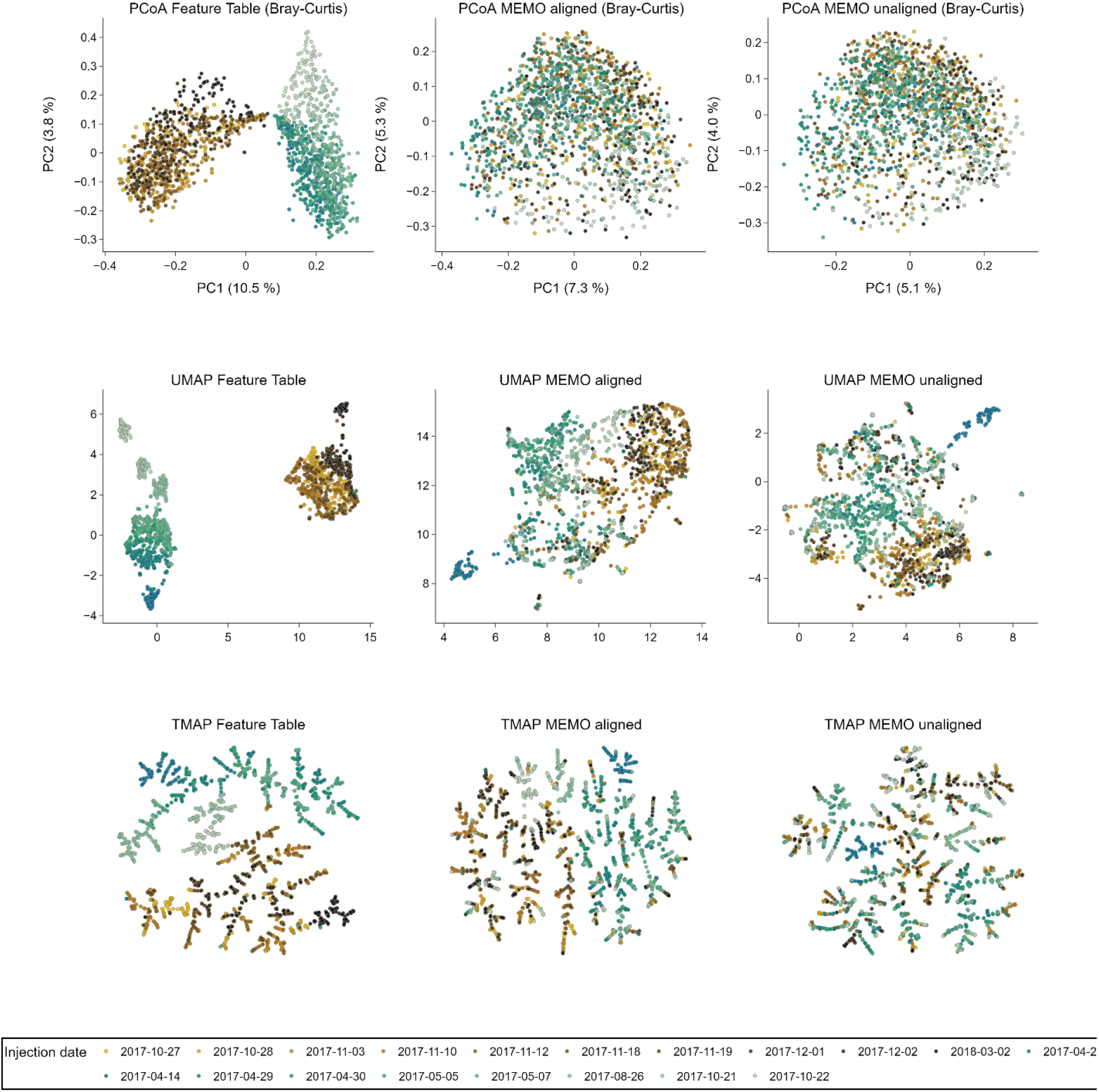
PCoA, UMAP and TMAP visualizations of the Feature Table, MEMO from aligned and MEMO from unaligned matrices of the plant extract dataset with samples colored according to their injection date.

**Figure S5:**
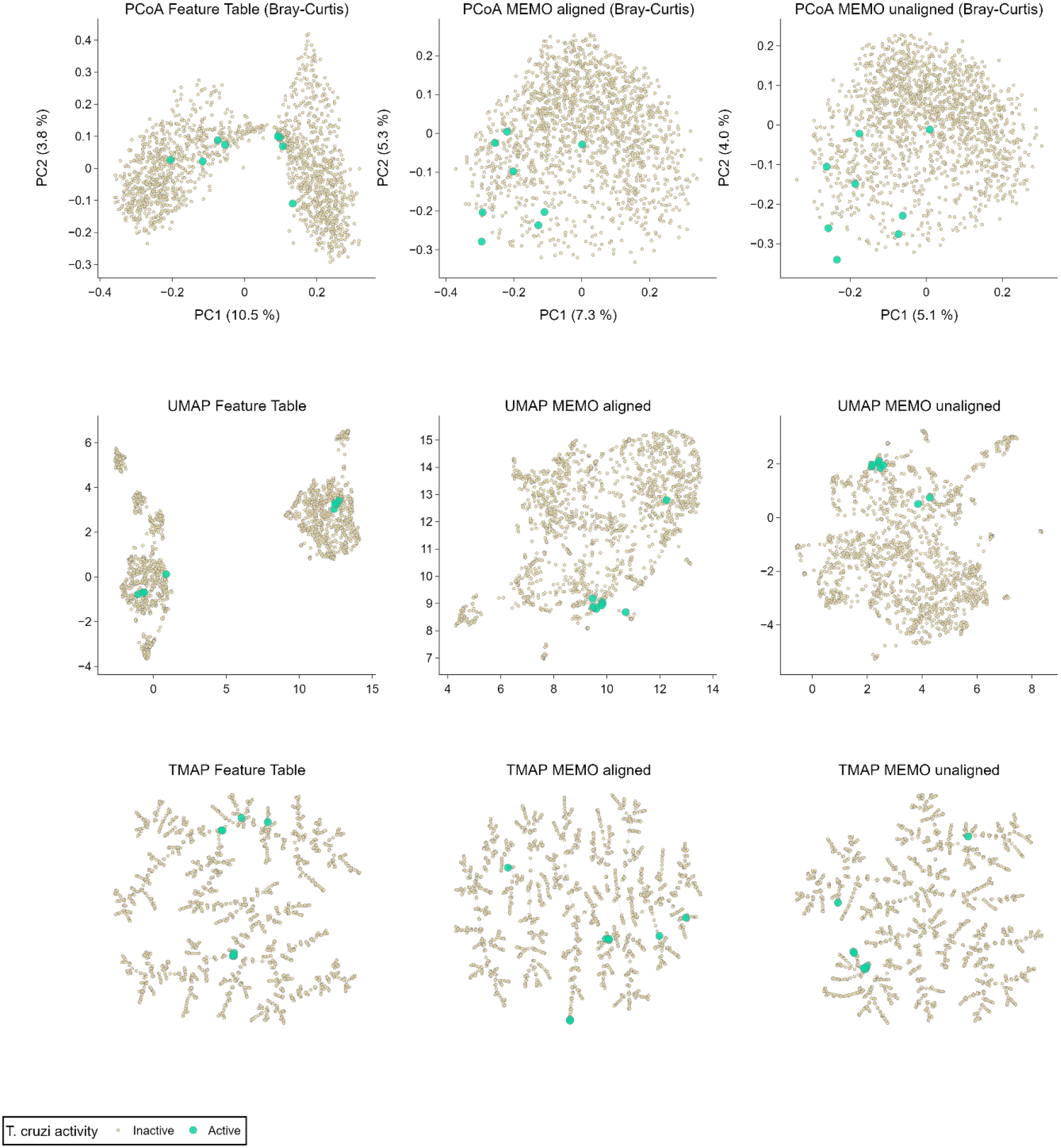
PCoA, UMAP and TMAP visualizations of the Feature Table, MEMO from aligned and MEMO from unaligned matrices of the plant extract dataset with samples colored according to their activity against *Trypanosoma cruzi*.

## References

Ahmed, S. A., Gogal, R. M., Jr, and Walsh, J. E. (1994). A new rapid and simple non-radioactive assay to monitor and determine the proliferation of lymphocytes: an alternative to [3H]thymidine incorporation assay. J. Immunol. Methods 170, 211–224.

Allard, P.-M., Péresse, T., Bisson, J., Gindro, K., Marcourt, L., Pham, V. C., et al. (2016). Integration of Molecular Networking and In-Silico MS/MS Fragmentation for Natural Products Dereplication. Anal. Chem. 88, 3317–3323.

Anderson, M. J. (2017). Permutational multivariate analysis of variance (PERMANOVA). Wiley StatsRef: Statistics Reference Online, 1–15. doi:10.1002/9781118445112.stat07841.

Arens, N., Döll, S., and Mock, H.-P. (2015). The reproducibility of liquid chromatography separation technology and its potential impact on large scale plant metabolomics experiments. J. Chromatogr. B Analyt. Technol. Biomed. Life Sci. 991, 41–45.

Bittremieux, W., Chen, C., Dorrestein, P. C., Schymanski, E. L., Schulze, T., Neumann, S., et al. (2020). Universal MS/MS Visualization and Retrieval with the Metabolomics Spectrum Resolver Web Service. bioRxiv, 2020.05.09.086066. doi:10.1101/2020.05.09.086066.

Buckner, F. S., Verlinde, C. L., La Flamme, A. C., and Van Voorhis, W. C. (1996). Efficient technique for screening drugs for activity against Trypanosoma cruzi using parasites expressing beta-galactosidase. Antimicrob. Agents Chemother. 40, 2592–2597.

Caporaso, J. G., Kuczynski, J., Stombaugh, J., Bittinger, K., Bushman, F. D., Costello, E. K., et al. (2010). QIIME allows analysis of high-throughput community sequencing data. Nat. Methods 7, 335–336.

Chambers, M. C., Maclean, B., Burke, R., Amodei, D., Ruderman, D. L., Neumann, S., et al. (2012). A cross-platform toolkit for mass spectrometry and proteomics. Nat. Biotechnol. 30, 918–920.

Cretton, S., Breant, L., Pourrez, L., Ambuehl, C., Marcourt, L., Ebrahimi, S. N., et al. (2014). Antitrypanosomal quinoline alkaloids from the roots of Waltheria indica. J. Nat. Prod. 77, 2304–2311.

Cretton, S., Bréant, L., Pourrez, L., Ambuehl, C., Perozzo, R., Marcourt, L., et al. (2015). Chemical constituents from Waltheria indica exert in vitro activity against Trypanosoma brucei and T. cruzi. Fitoterapia 105, 55–60.

Cretton, S., Dorsaz, S., Azzollini, A., Favre-Godal, Q., Marcourt, L., Ebrahimi, S. N., et al. (2016). Antifungal Quinoline Alkaloids from Waltheria indica. J. Nat. Prod. 79, 300–307.

Cretton, S., Kaiser, M., Karimou, S., Ebrahimi, S. N., Mäser, P., Cuendet, M., et al. (2020). Pyridine-4(1H)-one Alkaloids from Waltheria indica as Antitrypanosomatid Agents. J. Nat. Prod. 83, 3363–3371.

Duhrkop, K., Fleischauer, M., Ludwig, M., Aksenov, A. A., Melnik, A. V., Meusel, M., et al. (2019). SIRIUS 4: a rapid tool for turning tandem mass spectra into metabolite structure information. Nat. Methods 16, 299–302.

Duhrkop, K., Shen, H., Meusel, M., Rousu, J., and Bocker, S. (2015). Searching molecular structure databases with tandem mass spectra using CSI:FingerID. Proc. Natl. Acad. Sci. U. S. A. 112, 12580–12585.

Dunn, W. B., Wilson, I. D., Nicholls, A. W., and Broadhurst, D. (2012). The importance of experimental design and QC samples in large-scale and MS-driven untargeted metabolomic studies of humans. Bioanalysis 4, 2249–2264.

European Commission (2020). Official European Commission register of collections. Available at: https://ec.europa.eu/environment/nature/biodiversity/international/abs/pdf/Register%20of%20Collections.pdf.

Haug, K., Cochrane, K., Nainala, V. C., Williams, M., Chang, J., Jayaseelan, K. V., et al. (2020). MetaboLights: a resource evolving in response to the needs of its scientific community. Nucleic Acids Res. 48, D440–D444.

Huber, F., Ridder, L., Verhoeven, S., Spaaks, J. H., Diblen, F., Rogers, S., et al. (2021). Spec2Vec: Improved mass spectral similarity scoring through learning of structural relationships. PLoS Comput. Biol. 17, e1008724.

Huber, F., Verhoeven, S., Meijer, C., Spreeuw, H., Castilla, E., Geng, C., et al. (2020). Matchms - processing and similarity evaluation of mass spectrometry data. J. Open Source Softw. 5, 2411.

Jarmusch, A. K., Wang, M., Aceves, C. M., Advani, R. S., Aguirre, S., Aksenov, A. A., et al. (2020). ReDU: a framework to find and reanalyze public mass spectrometry data. Nat. Methods. doi:10.1038/s41592-020-0916-7.

Jarmusch, S. A., van der Hooft, J. J. J., Dorrestein, P. C., and Jarmusch, A. K. (2021). Advancements in capturing and mining mass spectrometry data are transforming natural products research. Nat. Prod. Rep. doi:10.1039/d1np00040c.

Junker, R. R. (2018). A biosynthetically informed distance measure to compare secondary metabolite profiles. Chemoecology 28, 29–37.

Leek, J. T., Scharpf, R. B., Bravo, H. C., Simcha, D., Langmead, B., Johnson, W. E., et al. (2010). Tackling the widespread and critical impact of batch effects in high-throughput data. Nat. Rev. Genet. 11, 733–739.

Lozupone, C., and Knight, R. (2005). UniFrac: a new phylogenetic method for comparing microbial communities. Appl. Environ. Microbiol. 71, 8228–8235.

Ludwig, M., Nothias, L.-F., Dührkop, K., Koester, I., Fleischauer, M., Hoffmann, M. A., et al. (2020). Database-independent molecular formula annotation using Gibbs sampling through ZODIAC. Nature Machine Intelligence 2, 629–641.

McInnes, L., Healy, J., and Melville, J. (2018). UMAP: Uniform Manifold Approximation and Projection for Dimension Reduction. arXiv [stat.ML]. Available at: http://arxiv.org/abs/1802.03426.

Myers, O. D., Sumner, S. J., Li, S., Barnes, S., and Du, X. (2017). One Step Forward for Reducing False Positive and False Negative Compound Identifications from Mass Spectrometry Metabolomics Data: New Algorithms for Constructing Extracted Ion Chromatograms and Detecting Chromatographic Peaks. Anal. Chem. 89, 8696–8703.

Nothias, L.-F., Petras, D., Schmid, R., Dührkop, K., Rainer, J., Sarvepalli, A., et al. (2020). Feature-based molecular networking in the GNPS analysis environment. Nat Methods. doi:10.1038/s41592-020-0933-6.

Page, B., Page, M., and Noel, C. (1993). A new fluorometric assay for cytotoxicity measurements in-vitro. Int. J. Oncol. 3, 473–476.

Pluskal, T., Castillo, S., Villar-Briones, A., and Orešič, M. (2010). MZmine 2: Modular framework for processing, visualizing, and analyzing mass spectrometry-based molecular profile data. BMC Bioinformatics 11, 395.

Probst, D., and Reymond, J.-L. (2020). Visualization of very large high-dimensional data sets as minimum spanning trees. J. Cheminform. 12. doi:10.1186/s13321-020-0416-x.

Rutz, A., Dounoue-Kubo, M., Ollivier, S., Bisson, J., Bagheri, M., Saesong, T., et al. (2019). Taxonomically Informed Scoring Enhances Confidence in Natural Products Annotation. Front. Plant Sci. 10. doi:10.3389/fpls.2019.01329.

Rutz, A., Sorokina, M., Galgonek, J., Mietchen, D., Willighagen, E., Gaudry, A., et al. (2021). The LOTUS Initiative for Open Natural Products Research: Knowledge Management through Wikidata. bioRxiv, 2021.02.28.433265. doi:10.1101/2021.02.28.433265.

Sedio, B. E., Rojas Echeverri, J. C., Boya P, C. A., and Wright, S. J. (2017). Sources of variation in foliar secondary chemistry in a tropical forest tree community. Ecology 98, 616–623.

Smith, C. A., Want, E. J., O’Maille, G., Abagyan, R., and Siuzdak, G. (2006). XCMS: processing mass spectrometry data for metabolite profiling using nonlinear peak alignment, matching, and identification. Anal. Chem. 78, 779–787.

Sumner, L. W., Amberg, A., Barrett, D., Beale, M. H., Beger, R., Daykin, C. A., et al. (2007). Proposed minimum reporting standards for chemical analysis Chemical Analysis Working Group (CAWG) Metabolomics Standards Initiative (MSI). Metabolomics 3, 211–221.

Tripathi, A., Vázquez-Baeza, Y., Gauglitz, J. M., Wang, M., Dührkop, K., Nothias-Esposito, M., et al. (2021). Chemically informed analyses of metabolomics mass spectrometry data with Qemistree. Nat. Chem. Biol. 17, 146–151.

Tsugawa, H., Cajka, T., Kind, T., Ma, Y., Higgins, B., Ikeda, K., et al. (2015). MS-DIAL: data-independent MS/MS deconvolution for comprehensive metabolome analysis. Nat. Methods 12, 523–526.

van der Hooft, J. J. J., Wandy, J., Barrett, M. P., Burgess, K. E. V., and Rogers, S. (2016). Topic modeling for untargeted substructure exploration in metabolomics. Proc. Natl. Acad. Sci. U. S. A. 113, 13738–13743.

Virtanen, P., Gommers, R., Oliphant, T. E., Haberland, M., Reddy, T., Cournapeau, D., et al. (2020). SciPy 1.0: fundamental algorithms for scientific computing in Python. Nat. Methods 17, 261–272.

Wang, M., Carver, J. J., Phelan, V. V., Sanchez, L. M., Garg, N., Peng, Y., et al. (2016). Sharing and community curation of mass spectrometry data with GNPS. Nat. Biotechnol. 34, 828–837.

Wehrens, R., Hageman, J. A., van Eeuwijk, F., Kooke, R., Flood, P. J., Wijnker, E., et al. (2016). Improved batch correction in untargeted MS-based metabolomics. Metabolomics 12, 88.

Wolfender, J. L., Litaudon, M., Touboul, D., and Queiroz, E. F. (2019). Innovative omics-based approaches for prioritisation and targeted isolation of natural products - new strategies for drug discovery. Nat. Prod. Rep. doi:10.1039/c9np00004f.

